# Decline of humoral responses against SARS-CoV-2 Spike in convalescent individuals

**DOI:** 10.1101/2020.07.09.194639

**Authors:** Guillaume Beaudoin-Bussières, Annemarie Laumaea, Sai Priya Anand, Jérémie Prévost, Romain Gasser, Guillaume Goyette, Halima Medjahed, Josée Perreault, Tony Tremblay, Antoine Lewin, Laurie Gokool, Chantal Morrisseau, Philippe Bégin, Cécile Tremblay, Valérie Martel-Laferrière, Daniel E. Kaufmann, Jonathan Richard, Renée Bazin, Andrés Finzi

**Author notes:** Contributed equally (Guillaume Beaudoin-Bussières, Annemarie Laumaea, Sai Priya Anand, Jérémie Prévost, Romain Gasser).

## Abstract

In the absence of effective vaccines and with limited therapeutic options, convalescent plasma is being collected across the globe for potential transfusion to COVID-19 patients. The therapy has been deemed safe and several clinical trials assessing its efficacy are ongoing. While it remains to be formally proven, the presence of neutralizing antibodies is thought to play a positive role in the efficacy of this treatment. Indeed, neutralizing titers of ≥1:160 have been recommended in some convalescent plasma trials for inclusion. Here we performed repeated analyses at one-month interval on 31 convalescent individuals to evaluate how the humoral responses against the SARS-CoV-2 Spike, including neutralization, evolve over time. We observed that receptor-binding domain (RBD)-specific IgG slightly decreased between six and ten weeks after symptoms onset but RBD-specific IgM decreased much more abruptly. Similarly, we observed a significant decrease in the capacity of convalescent plasma to neutralize pseudoparticles bearing SARS-CoV-2 S wild-type or its D614G variant. If neutralization activity proves to be an important factor in the clinical efficacy of convalescent plasma transfer, our results suggest that plasma from convalescent donors should be recovered rapidly after symptoms resolution.

## MAIN TEXT

Until an efficient vaccine to protect from SARS-CoV-2 infection is available, alternative approaches to treat or prevent acute COVID-19 are urgently needed. A promising approach is the use of convalescent plasma containing anti-SARS-CoV-2 antibodies collected from donors who have recovered from COVID-19 (1). Convalescent plasma therapy was successfully used in the treatment of SARS, MERS and influenza H1N1 pandemics and was associated with improvement of clinical outcomes (2–4). Experience to date shows that the passive transfer of convalescent plasma to acute COVID-19 patients has been shown to be well tolerated and presented some hopeful signs (5–9). In one study, the convalescent plasma used had high titers of IgG to SARS-CoV-2 (at least 1:1640), which correlated positively with neutralizing activity (10). While it remains to be formally demonstrated, neutralizing activity is considered an important determinant of convalescent plasma efficacy (11) and regulatory agencies have been recommending specific thresholds for qualifying convalescent plasma prior to its release. While neutralizing function has been associated with protection against reinfection in rhesus macaques (12), other antibody functions may be relevant for controlling an acute infection and should be examined to better understand the correlates of convalescent plasma-mediated efficacy (7).

It was recently reported that the humoral responses against SARS-CoV-2 are built rapidly, peaking at weeks 2 or 3 after symptoms onset but steadily decreases thereafter (13–15). Moreover, in a cross-sectional study we reported that the neutralization capacity decreased between the third and the sixth week after symptoms onset (14). Since convalescent patients are generally required to wait for 14 days after recovery to start plasma donations and that they may give multiple times in the following weeks, most donations are likely to occur even later than this. Whether the neutralization capacity of convalescent plasma is stabilized after six weeks or decreases further remains unknown. To answer this question, which might have practical implications in the selection of plasma from convalescent donors, we analyzed serological samples from 31 convalescent donors collected at six and ten weeks after symptoms onset.

All convalescent donors initially tested positive for SARS-CoV-2 by RT-PCR on nasopharyngeal specimens with complete resolution of symptoms for at least 14 days before blood sampling. The average age of the donors was 46 years old (22 males and 9 females). We collected plasma samples from each individual at two time-points: 6 weeks after symptoms onset (baseline, median 43 days) and 4 weeks after (1 month, median 74 days after symptoms onset) (Supplemental Table 1).

We first evaluated the presence of RBD-specific IgG and IgM antibodies by ELISA as we recently described (14). In agreement with a recent report (16), we observed that both RBD-specific IgG and IgM titers significantly decreased between 6 and 10 weeks after symptoms onset. We noted that IgM titers diminished significantly more abruptly than IgG titers (Figure 1B and E respectively). Accordingly, the percentage of convalescent individuals presenting detectable titers of IgM decreased by ~13% at 10 weeks after symptoms onset (Figure 1C) while the percentage of infected individuals presenting detectable titers of IgG remained stable (Figure 1F).

**Figure 1.**
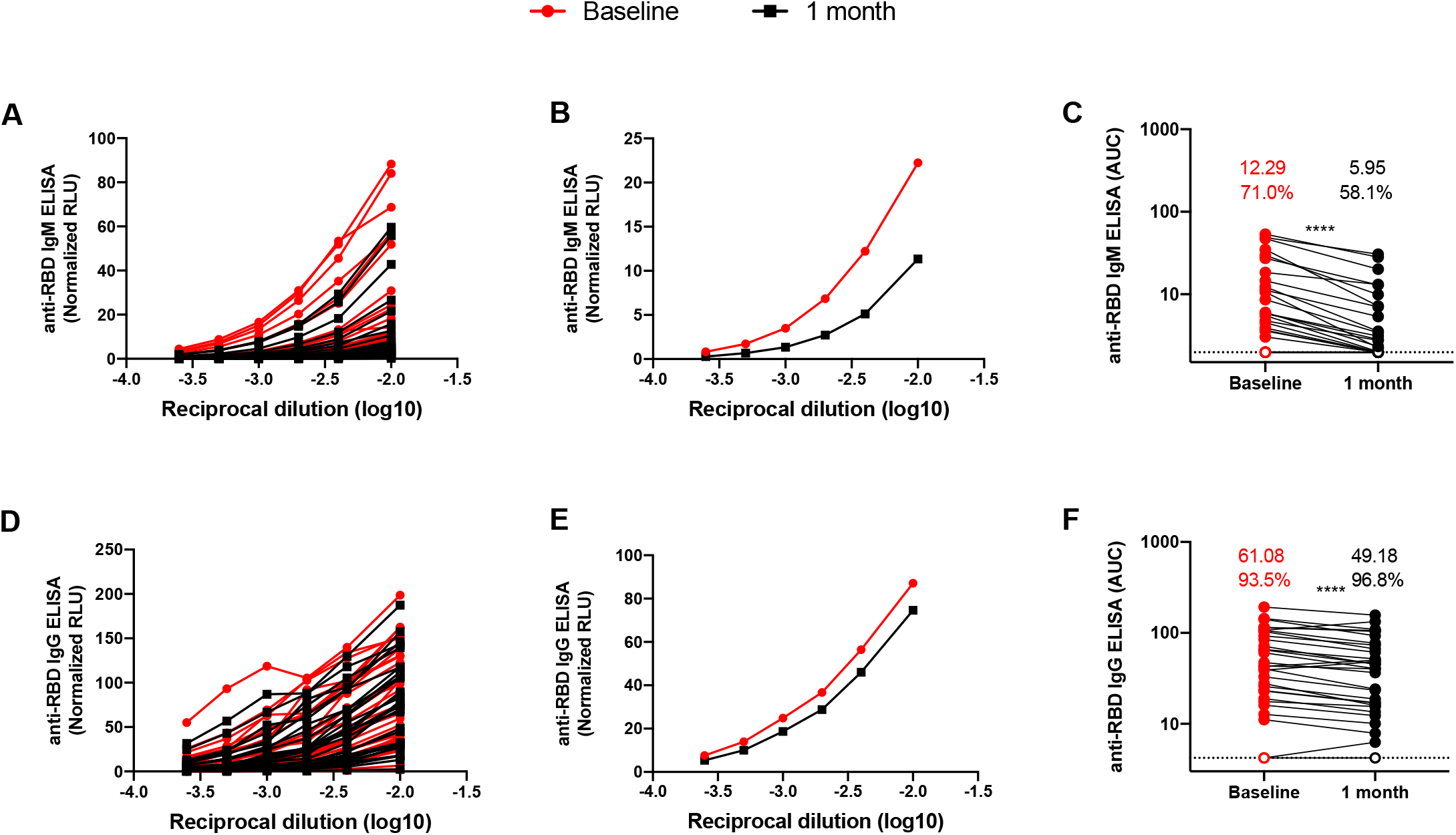
SARS-CoV-2 RBD-specific IgM and IgG decrease over time. Indirect ELISA was performed using recombinant SARS-CoV-2 RBD and incubated with plasma samples recovered at baseline (6 weeks after symptoms onset; red lines) and 1 month later (black lines). Anti-RBD antibody binding was detected using (A-C) anti-IgM-HRP or (D-F) anti-IgG-HRP. Relative light units (RLU) obtained with BSA (negative control) were subtracted and further normalized to the signal obtained with the anti-RBD CR3022 mAb present in each plate. Data in graphs represent RLU (A,D) done in triplicate for each plasma or (B, E) the mean of the plasma recovered at baseline (red) and 1 month later (black). (C, F) Areas under the curve (AUC) were calculated based on RLU datasets shown in (A, D) using GraphPad Prism software and their average is shown on top of panels C and F, the percentage (%) of samples presenting a positive signal is indicated. Undetectable measures are represented as white symbols and limits of detection are plotted (calculated with samples from COVID-19 negative patients). Statistical significance was tested using Wilcoxon matched-pairs signed rank test (**** p < 0.0001).

We next used flow cytometry to examine the ability of convalescent plasma to recognize the full-length SARS-CoV-2 Spike expressed at the cell surface. Briefly, 293T cells expressing SARS-CoV-2 S glycoproteins were stained with plasma samples, followed by incubation with secondary antibodies recognizing all antibody isotypes. Since the SARS-CoV-2 strain circulating in Europe and North America has the D614G mutation (17), we also evaluated recognition of this variant by flow cytometry. As presented in Supplemental Figure 1A, convalescent plasma from 96.8 % of donors (all but one) recognized both SARS-CoV-2 S (WT and D614G) at baseline. While this percentage remained stable four weeks later, the recognition (mean fluorescence intensity, MFI) was significantly diminished for both WT and D614G S-expressing cells, indicating that Spike-reactive antibodies were less abundant in convalescent plasma collected at this later time point. Interestingly, the MFI were almost identical for cells expressing the WT or D614G variant S (7206 and 7209 respectively, Fig. S1A), suggesting that the mutation did not significantly affect S conformation. In agreement with recent work, we observed that SARS-CoV-2-elicited antibodies cross-react with human *Sarbecoviruses* (14) (SARS-CoV; Figure S1B) and with another *Betacoronavirus* (OC43) whereas no cross-reactive antibodies to *Alphacoronavirus* (NL63, 229E) S glycoproteins (Figure S1C and S2) were detected. Cross-reactive antibodies recognizing SARS-CoV and OC43 S glycoproteins decreased between the two time-points following a trend similar to SARS-CoV-2 S-reactive antibodies.

We next measured the capacity of plasma samples to neutralize pseudoparticles bearing SARS-CoV-2 S, its D614G variant, SARS-CoV S or VSV-G glycoproteins using 293T cells stably expressing ACE2 as target cells (Figure 2). Neutralizing activity against SARS-CoV-2 WT or D614G S glycoprotein, as measured by the neutralization half-maximum inhibitory dilution (ID50), was detected in 71% of patients six weeks after symptoms onset. SARS-CoV-2 neutralization was specific since no neutralization was observed against pseudoparticles expressing VSV-G. Neutralizing activity against pseudoparticles bearing the SARS-CoV S glycoprotein was detected in only 25% of convalescent plasma and exhibited low potency, as previously reported (Figure 2) (14). Of note, while we observed enhanced infectivity for the D614G variant compared to its WT SARS-CoV-2 S counterpart (Figure S3A), no major differences in neutralization with convalescent plasma were detected at both time-points (Figure S3B), thus suggesting that the D614G change does not affect the overall conformation of the Spike, in agreement with recent findings (18).

**Figure 2.**
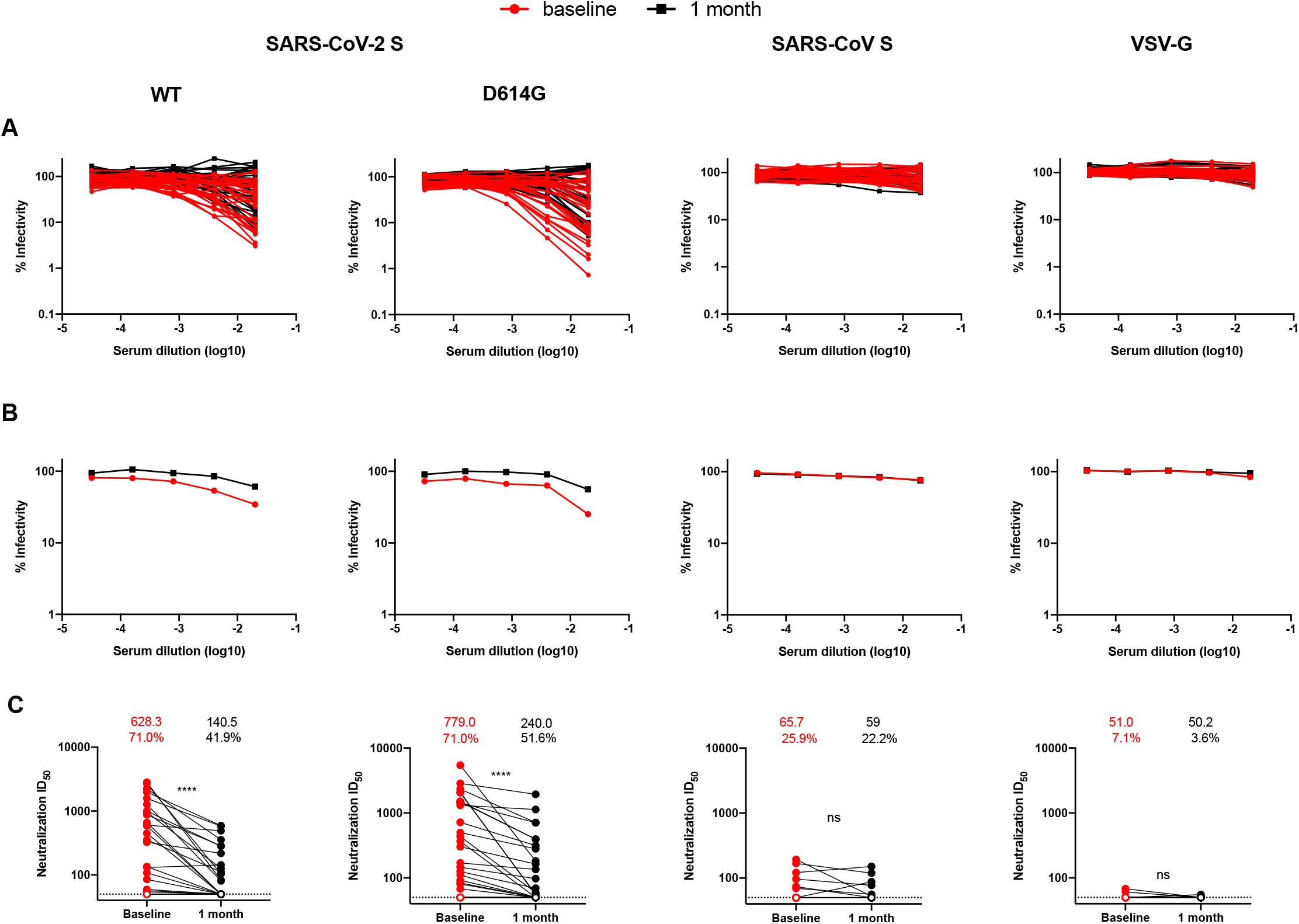
Neutralizing activity of convalescent plasma decreases over time. (A) Pseudoviral particles coding for the luciferase reporter gene and bearing the following glycoproteins: SARS-CoV-2 S or its D614G counterpart, SARS-CoV S and VSV-G were used to infect 293T-ACE2 cells. Pseudoviruses were incubated with serial dilutions of plasma samples recovered at baseline (6 weeks after symptoms onset) or collected 1 month later, at 37°C for 1h prior to infection of 293T-ACE2 cells. Infectivity at each dilution was assessed in duplicate and is shown as the percentage of infection without plasma for each pseudovirus. (B) The median of neutralization by baseline (red) or 1 month (black) plasma samples is shown. (C) Neutralization half maximal inhibitory serum dilution (ID_50_) values were determined using a normalized non-linear regression with Graphpad Prism software. Undetectable measures measures (ID_50_< 50) are represented as white symbols. The mean neutralizing titers and the proportion (%) of neutralizer (patients with an ID50 over 50) are shown above the graphs. Statistical significance was tested using Wilcoxon matched-pairs signed rank tests (ns, not significant; **** p < 0.0001).

The capacity to neutralize SARS-CoV-2 S WT or D614G-pseudotyped particles significantly correlated with the presence of RBD-specific IgG, IgM and anti-S antibodies (Figure S4). Interestingly, we observed a pronounced decrease (20-30%) in the percentage of patients able to neutralize pseudoparticles bearing SARS-CoV-2 S glycoprotein between 6 and 10 weeks after symptoms onset. Moreover, with plasma that still neutralized, the neutralization activity significantly decreased between these two time-points (Figure 2C). Interestingly, RBD-specific IgM and neutralizing activity declined more significantly in convalescent plasma overtime compared to RBD-specific IgG and anti-S Abs (Figure S5A, B). Moreover, while the loss of neutralizing activity on the WT and D614G pseudoparticles over time correlated with the loss of anti-RBD IgM and IgG antibodies, the correlation was higher for IgM than IgG (Figure S5C, D), suggesting that at least part of the neutralizing activity could be mediated by IgM, as recently proposed (13, 14).

In summary, our study indicates that plasma neutralization activity keeps decreasing passed the sixth week of symptom onset (14). It is currently unknown whether neutralizing activity is truly driving the efficacy of convalescent plasma in acute COVID-19. If this was found to be the case, our results suggest that efforts should be made to ensure convalescent plasma is collected as soon as possible after recovery from active infection.

## ACKNOWLEDGMENTS

The authors thank the convalescent plasma donors who participated in this study, the Héma-Québec team involved in convalescent donor recruitment and plasma collection, the CRCHUM BSL3 and Flow Cytometry Platforms for technical assistance, Dr Stefan Pöhlmann (Georg-August University, Germany) for the plasmids coding for SARS-CoV-2 S, 229E S and NL63 S glycoproteins, Dr Marcelline Côté (University of Ottawa) for the OC43 S expressor and Dr M. Gordon Joyce (U.S. MHRP) for the monoclonal antibody CR3022. This work was supported by le Ministère de l’Économie et de l’Innovation du Québec, Programme de soutien aux organismes de recherche et d’innovation to A.F and by the Fondation du CHUM. This work was also supported by a CIHR foundation grant #352417 to A.F. A.F. is the recipient of a Canada Research Chair on Retroviral Entry # RCHS0235 950-232424. G.B.B., S.P.A and J.P. are supported by CIHR fellowships. R.G. is supported by a MITACS Accélération postdoctoral fellowship. V.M.L. and P.B. are supported by FRQS Junior 1 salary awards. D.E.K. is a FRQS Merit Research Scholar. The funders had no role in study design, data collection and analysis, decision to publish, or preparation of the manuscript. The authors declare no competing interests.

## SUPPLEMENTAL MATERIAL

TEXT S1

FIG S1

FIG S2

FIG S3

FIG S4

FIG S5

TABLE S1

## SUPPLEMENTAL MATERIAL AND METHODS

### Ethics statement

All work was conducted in accordance with the Declaration of Helsinki in terms of informed consent and approval by an appropriate institutional board. Convalescent plasmas were obtained from donors who consented to participate in this research project at Héma-Québec (REB # 2020-004) and CHUM (19.381). The donors met all donor eligibility criteria: previous confirmed COVID-19 infection and complete resolution of symptoms for at least 14 days.

### Plasmids

The plasmids expressing the human coronavirus Spikes of SARS-CoV-2, SARS-CoV, NL63 229E (1, 2) and OC43 (3) were previously described. The D614G mutation was introduced using the QuikChange II XL site-directed mutagenesis protocol (Stratagene). The presence of the desired mutations was determined by automated DNA sequencing. The plasmid encoding for SARS-CoV-2 S RBD was synthesized commercially by Genscript. The RBD sequence (encoding for residues 319-541) fused to a C-terminal hexahistidine tag was cloned into the pcDNA3.1(+) expression vector. The vesicular stomatitis virus G (VSV-G)-encoding plasmid (pSVCMV-IN-VSV-G) was previously described (4).

### Cell lines

293T human embryonic kidney cells (obtained from ATCC) were maintained at 37°C under 5% CO2 in Dulbecco’s modified Eagle’s medium (DMEM) (Wisent) containing 5% fetal bovine serum (VWR), 100 UI/ml of penicillin and 100μg/ml of streptomycin (Wisent). 293T-ACE2 cell line was previously reported (3).

### Protein expression and purification

FreeStyle 293F cells (Invitrogen) were grown in FreeStyle 293F medium (Invitrogen) to a density of 1 x 10^6^ cells/mL at 37°C with 8 % CO2 with regular agitation (150 rpm). Cells were transfected with a plasmid coding for SARS-CoV-2 S RBD using ExpiFectamine 293 transfection reagent, as directed by the manufacturer (Invitrogen). One week later, cells were pelleted and discarded. Supernatants were filtered using a 0.22 μm filter (Thermo Fisher Scientific). The recombinant RBD proteins were purified by nickel affinity columns, as directed by the manufacturer (Invitrogen). The RBD preparations were dialyzed against phosphate-buffered saline (PBS) and stored in aliquots at −80°C until further use. To assess purity, recombinant proteins were loaded on SDS-PAGE gels and stained with Coomassie Blue.

### Plasma and antibodies

Plasma from SARS-CoV-2-infected and uninfected donors were collected, heat-inactivated for 1 hour at 56 °C and stored at −80°C until ready to use in subsequent experiments. Plasma from uninfected donors were used as negative controls and used to calculate the seropositivity threshold in our ELISA and flow cytometry assays. The monoclonal antibody CR3022 was used as a positive control in ELISA assays and was previously described (5–7). Horseradish peroxidase (HRP)-conjugated antibody specific for the Fc region of human IgG (Invitrogen) or for the Fc region of human IgM (Jackson ImmunoResearch Laboratories, inc.) were used as secondary antibodies to detect antibody binding in ELISA experiments. Alexa Fluor-647-conjugated goat anti-human IgG (H+L) Abs (Invitrogen) were used as secondary antibodies to detect sera binding in flow cytometry experiments.

### ELISA

The SARS-CoV-2 RBD assay used was recently described (3). Briefly, recombinant SARS-CoV-2 S RBD proteins (2.5 μg/ml), or bovine serum albumin (BSA) (2.5 μg/ml) as a negative control, were prepared in PBS and were adsorbed to plates (MaxiSorp; Nunc) overnight at 4°C. Coated wells were subsequently blocked with blocking buffer (Tris-buffered saline [TBS] containing 0.1% Tween20 and 2% BSA) for 1h at room temperature. Wells were then washed four times with washing buffer (Tris-buffered saline [TBS] containing 0.1% Tween20). CR3022 mAb (50ng/ml) or serial dilutions of plasma from SARS-CoV-2-infected or uninfected donors (1/100; 1/250; 1/500; 1/1000; 1/2000; 1/4000) were prepared in a diluted solution of blocking buffer (0.1 % BSA) and incubated with the RBD-coated wells for 90 minutes at room temperature. Plates were washed four times with washing buffer followed by incubation with secondary Abs (diluted in a diluted solution of blocking buffer (0.4% BSA)) for 1h at room temperature, followed by four washes. HRP enzyme activity was determined after the addition of a 1:1 mix of Western Lightning oxidizing and luminol reagents (Perkin Elmer Life Sciences). Light emission was measured with a LB941 TriStar luminometer (Berthold Technologies). Signal obtained with BSA was subtracted for each plasma and was then normalized to the signal obtained with CR3022 mAb present in each plate. The seropositivity threshold was established using the following formula: mean of all COVID-19 negative plasmas + (3 standard deviation of the mean of all COVID-19 negative plasmas).

### Flow cytometry analysis of cell-surface staining

Using the standard calcium phosphate method, 10μg of Spike expressor and 2μg of a green fluorescent protein (GFP) expressor (pIRES-GFP) were transfected into 2 × 10^6^ 293T cells. At 48h post transfection, 293T cells were stained with plasma from SARS-CoV-2-infected or uninfected individuals (1:250 dilution). The percentage of transfected cells (GFP+ cells) was determined by gating the living cell population based on the basis of viability dye staining (Aqua Vivid, Invitrogen). Samples were acquired on a LSRII cytometer (BD Biosciences, Mississauga, ON, Canada) and data analysis was performed using FlowJo vX.0.7 (Tree Star, Ashland, OR, USA). The seropositivity threshold was established using the following formula: mean of all COVID-19 negative plasmas + (3 standard deviation of the mean of all COVID-19 negative plasma + inter-assay coefficient of variability).

### Virus neutralization assay

293T-ACE2 target cells were infected with single-round luciferase-expressing lentiviral particles. Briefly, 293T cells were transfected by the calcium phosphate method with the lentiviral vector pNL4.3 R-E-Luc (NIH AIDS Reagent Program) and a plasmid encoding for SARS-CoV-2 Spike (WT or D614G), SARS-CoV Spike or VSV-G at a ratio of 5:4. Two days post-transfection, cell supernatants were harvested and stored at –80°C until use. 293T-ACE2 target cells were seeded at a density of 1×10^4^ cells/well in 96-well luminometer-compatible tissue culture plates (Perkin Elmer) 24h before infection. Recombinant viruses in a final volume of 100μl were incubated with the indicated plasma dilutions (1/50; 1/250; 1/1250; 1/6250; 1/31250) for 1h at 37°C and were then added to the target cells followed by incubation for 48h at 37°C; cells were lysed by the addition of 30μl of passive lysis buffer (Promega) followed by one freeze-thaw cycle. An LB941 TriStar luminometer (Berthold Technologies) was used to measure the luciferase activity of each well after the addition of 100μl of luciferin buffer (15mM MgSO4, 15mM KPO4 [pH 7.8], 1mM ATP, and 1mM dithiothreitol) and 50μl of 1mM d-luciferin potassium salt (Prolume). The neutralization half-maximal inhibitory dilution (ID50) represents the plasma dilution to inhibit 50% of the infection of 293T-ACE2 cells by recombinant viruses bearing the indicated surface glycoproteins.

### Statistical analyses

Statistics were analyzed using GraphPad Prism version 8.0.2 (GraphPad, San Diego, CA, (USA). Every data set was tested for statistical normality and this information was used to apply the appropriate (parametric or nonparametric) statistical test. P values <0.05 were considered significant; significance values are indicated as * P<0.05, ** P<0.01, *** P<0.001, **** P<0.0001.

---

**Supplemental Table 1.** Cohort characteristics.

| Time after<br>symptoms onset<br>and first sample<br>collection :<br>Baseline<br>(Median; Days;<br>Day range) | Time after<br>symptoms onset<br>and second<br>sample<br>collection : 1<br>month (Median;<br>Days; Day range) | Age (Average;<br>Years; Age<br>range) | Sex |  |
| --- | --- | --- | --- | --- |
|  |  |  | Male (n) | Female (n) |
| 43 (16-60) | 74 (44-87) | 46 (20-67) | 22 | 9 |

**Figure S1.**
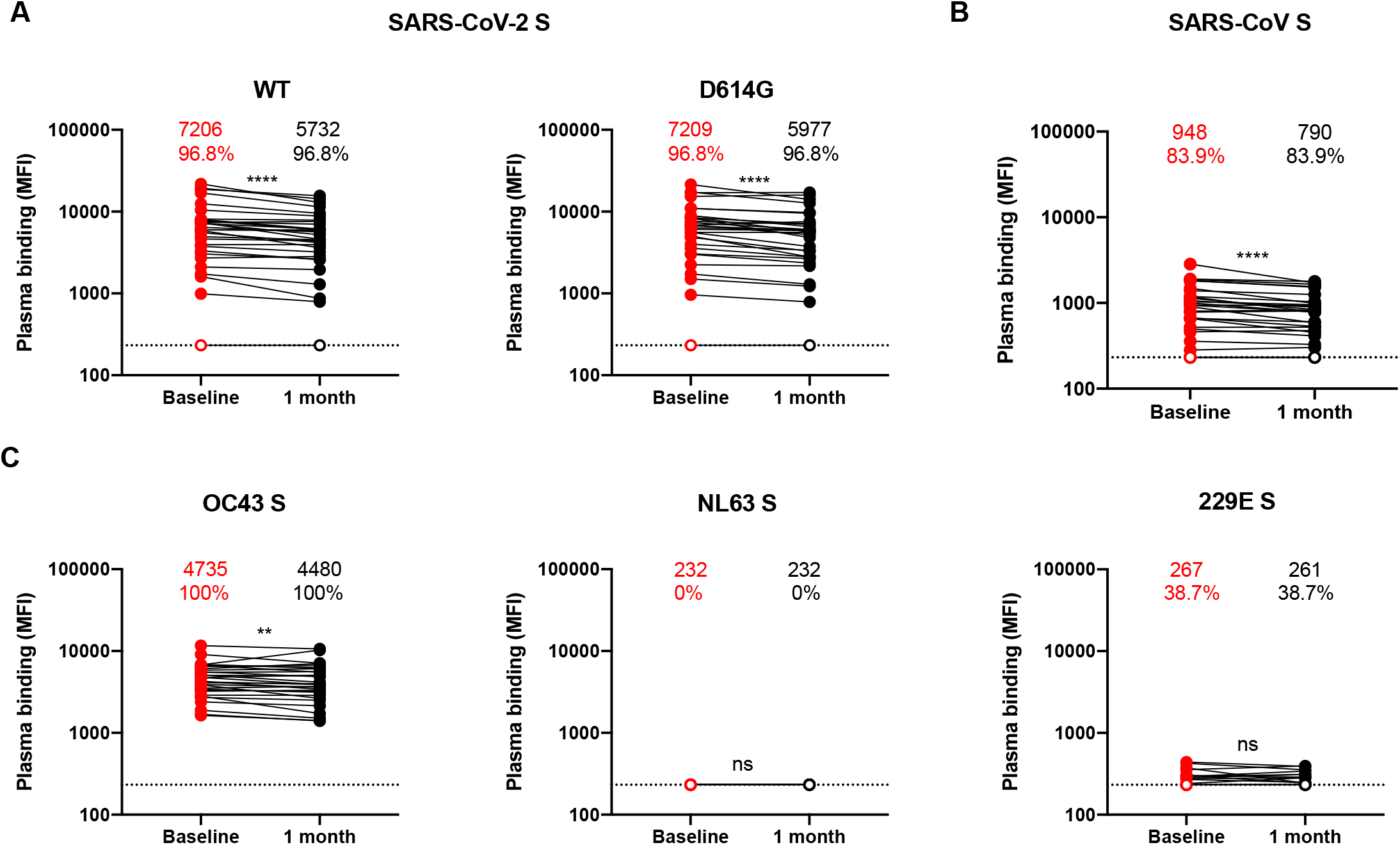
Cross-reactive antibodies against human *Betacoronaviruses* decrease over time. Cell-surface staining of 293T cells expressing full-length Spike (S) from different HCoV (A) SARS-CoV-2 or its D614G counterpart (B), SARS-CoV, (C) OC43, NL63 and 229E with plasma samples recovered at baseline (6 weeks after symptoms onset) and 1 month later. The graphs shown represent the median fluorescence intensities (MFI). Undetectable measures are represented as white symbols and limits of detection are plotted. The average MFI and percentage (%) of positive samples is indicated in top of each panel. Statistical significance was tested using Wilcoxon matched-pairs signed rank test (ns, not significant; ** p < 0.01; **** p < 0.0001).

**Figure S2.**
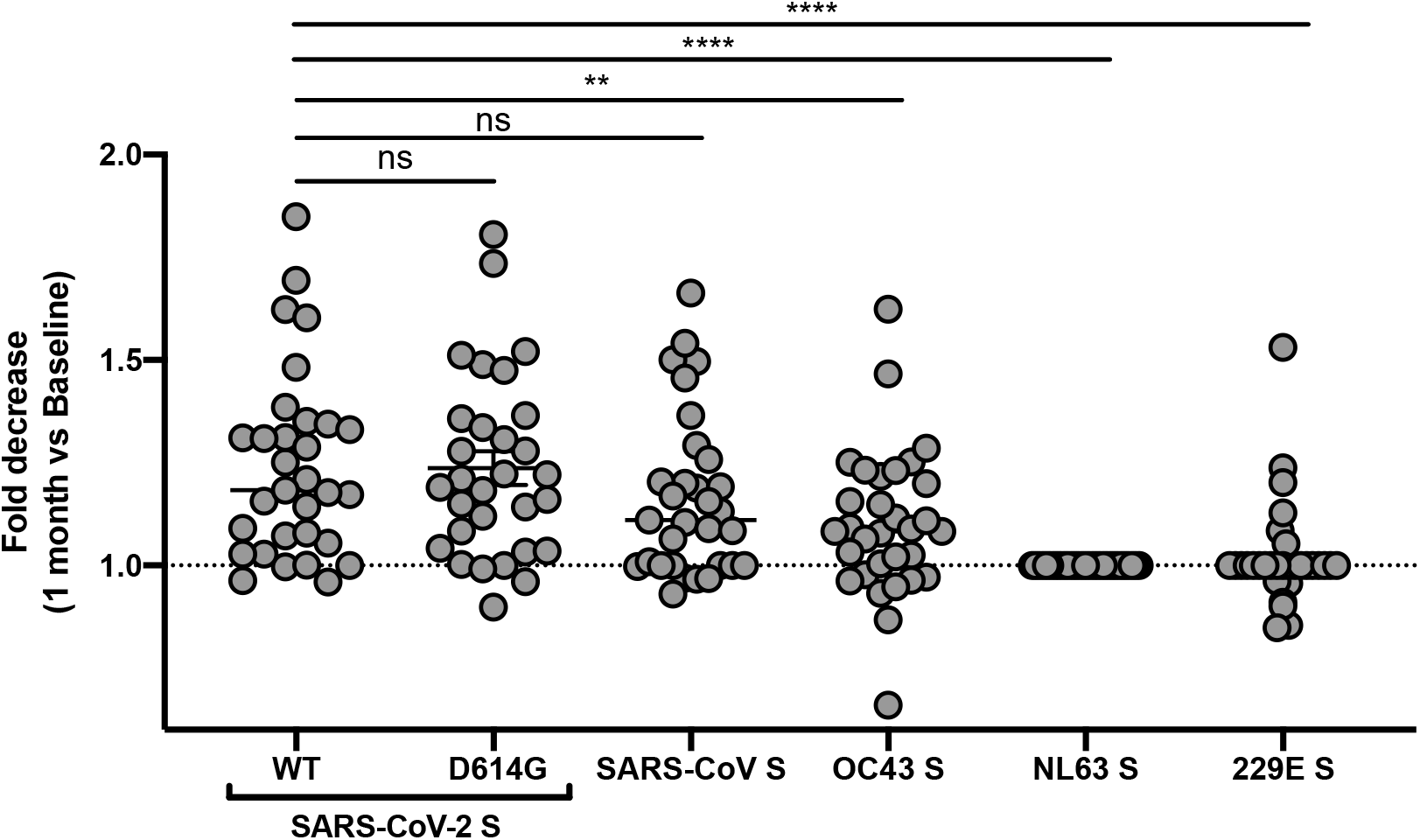
Decrease in cross-reactive antibodies. Fold decrease (1 month vs baseline) of the capacity of plasma to recognize by flow cytometry SARS-CoV-2 S WT, SARS-CoV-2 S D614G, SARS-CoV S, OC43 S, NL63 S and 229E S glycoproteins expressed at the surface of 293T cells. Statistical significance was tested using Wilcoxon matched-pairs signed rank tests (ns, not significant; ** p < 0.01; **** p < 0.0001).

**Figure S3.**
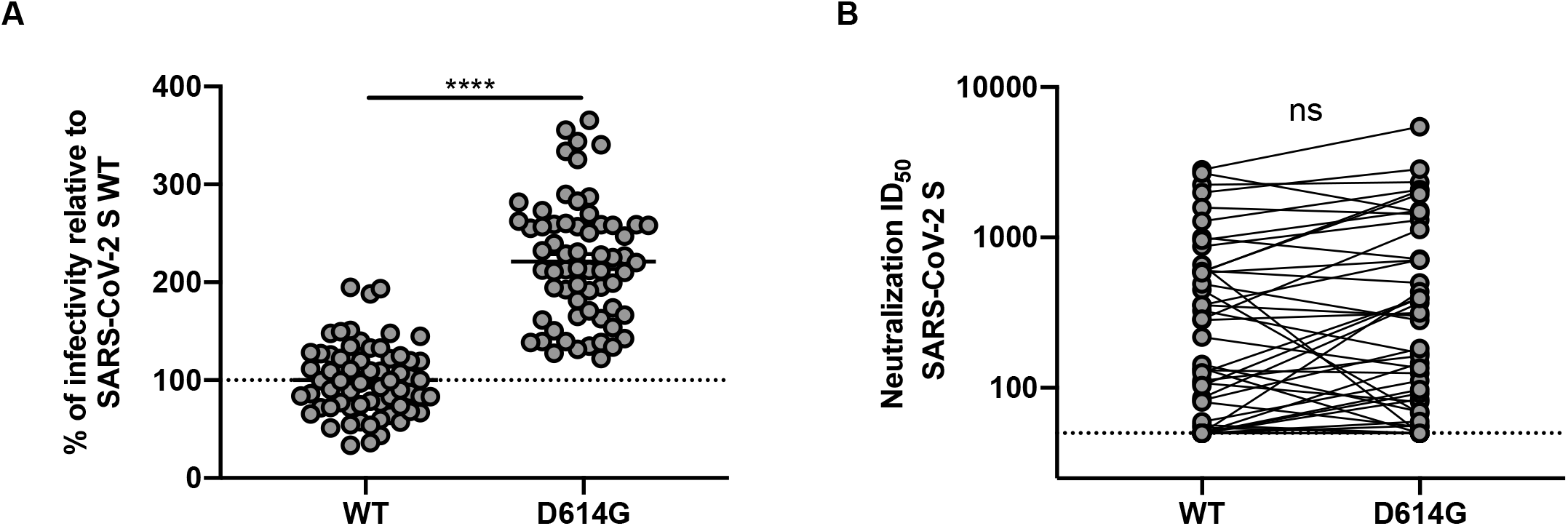
D614G mutation enhances SARS-CoV-2 infectivity but does not affect its susceptibility to plasma neutralization. (A) Reverse Transcriptase normalized levels of pseudoviral particles bearing the SARS-CoV-2 S WT or D614G variant were used to infect 293T/ACE2 cells and infectivity measured 48h later by luciferase activity. Graph shown represents the percentage of infectivity relative to pseudoviral particle bearing the SARS-CoV-2 S WT. Statistical significance was tested using Mann-Whitney U tests (**** p < 0.0001). (B) Comparison between the neutralization ID_50_ from pseudoparticles bearing SARS-CoV-2 S WT and SARS-CoV-2 S D614G. Statistical significance was tested using Wilcoxon matched-pairs signed rank test. (ns, not significant).

**Figure S4.**
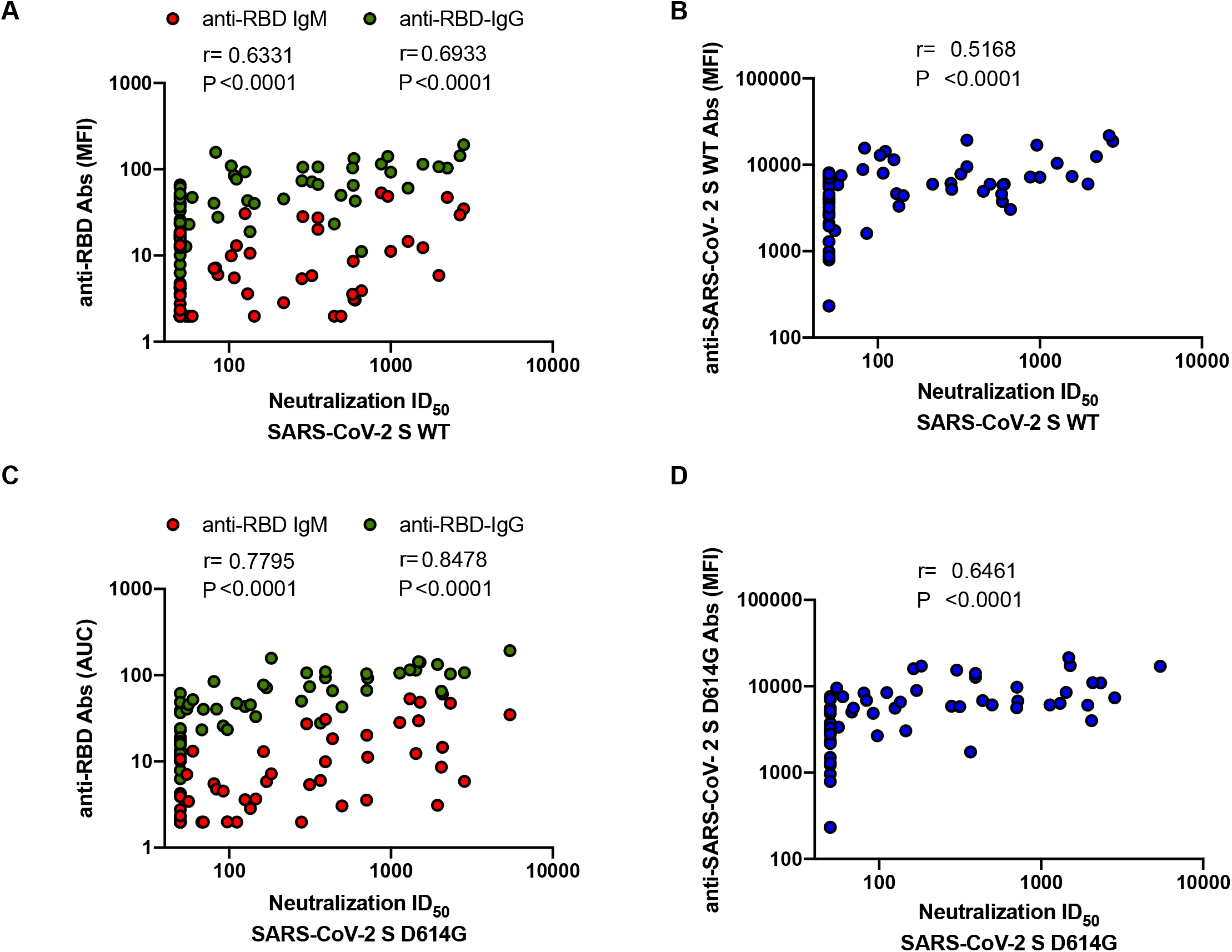
SARS-CoV-2 RBD and full length S specific antibodies correlate with pseudoviruses neutralization. Anti-RBD IgG and IgM evaluated by ELISA (A, C) or anti-S antibodies evaluated by flow cytometry (B, D) were plotted against the levels of neutralization (ID_50_) of pseudoparticles bearing the SARS-CoV-2 S (A, B) or its D614G counterpart (C, D). Statistical analysis was performed using Spearman rank correlation tests.

**Figure S5.**
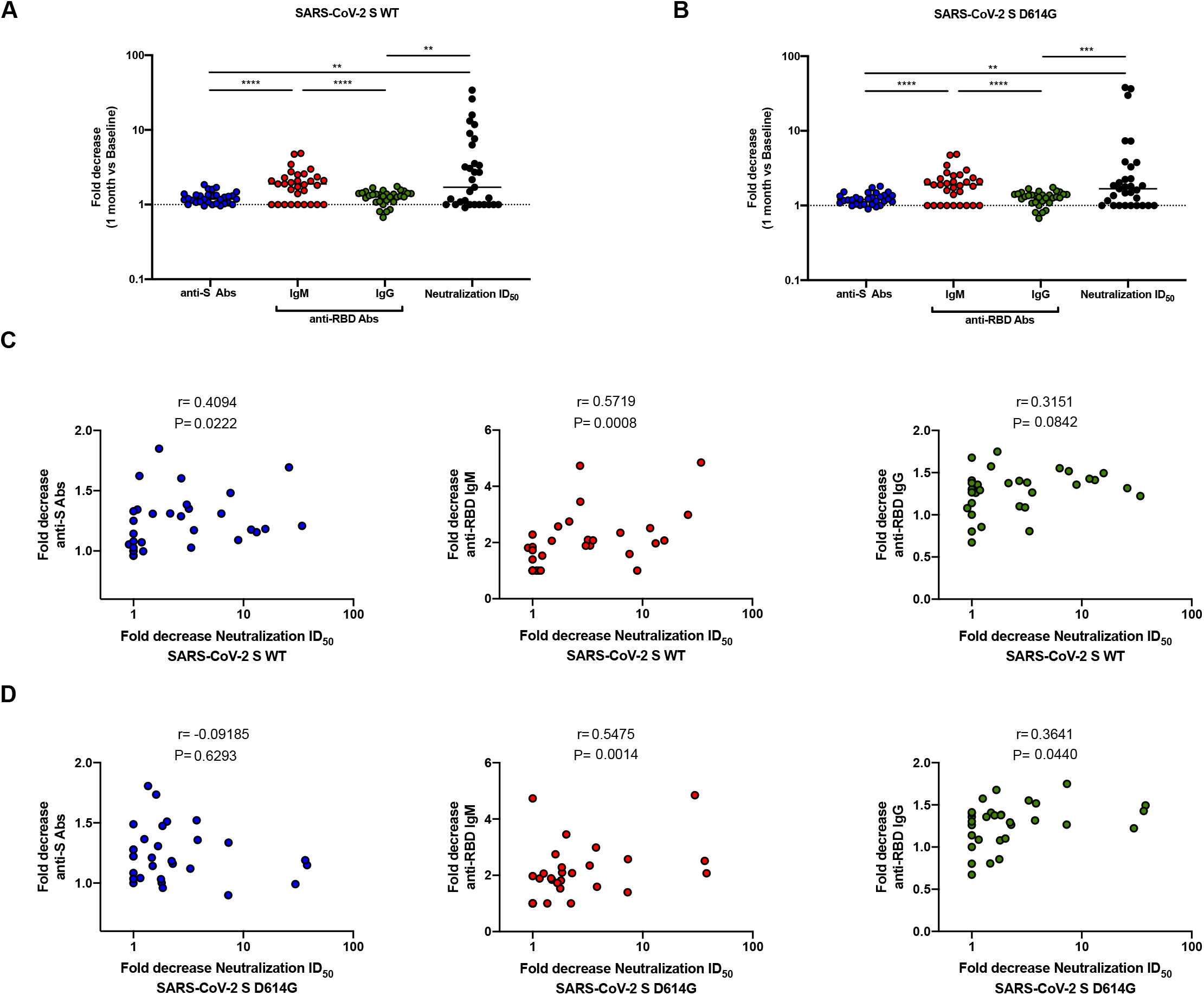
Decrease in anti-RBD IgM antibodies over time correlates with reduced neutralizing activity. Fold decrease of the 31 pairs of plasma over the course of 1 month (1 month over Baseline) of the levels of anti-SARS-CoV-2 S WT or D614G antibodies quantified by flow cytometry, anti-RBD antibodies (IgM and IgG) quantified by ELISA and of neutralization ID_50_ with pseudoparticules bearing (A) SARS-CoV-2 S WT or (B) SARS-CoV-2 S D614G. Correlation between the fold decrease over the course of 1 month of anti-SARS-CoV-2 S WT or D614G antibodies quantified by flow cytometry, anti-RBD (IgM and IgG) antibodies quantified by ELISA and the fold decrease of the neutralization ID_50_ of pseudoparticules bearing (C) SARS-CoV-2 S WT or (D) SARS-CoV-2 S D614G. (A, B) Statistical significance was tested using Wilcoxon matched-pairs signed rank tests (** p < 0.01; *** p < 0.001; **** p < 0.0001). (C, D) Statistical significance was tested using Spearman rank correlation tests.

